# Grey matter microstructural alterations distinguish cognitive impairment in Lewy body diseases

**DOI:** 10.64898/2026.01.06.697730

**Authors:** Angeliki Zarkali, Ivelina Dobreva, Naomi Hannaway, Rohan Bhome, George EC Thomas, Steffani Carrera, Amanda J Heslegrave, Elena Veleva, Henrik Zetterberg, Rimona S Weil

## Abstract

**INTRODUCTION.:** Despite widespread cortical involvement in Lewy body diseases, conventional grey matter MRI measures have limited sensitivity. Microstructural measures from diffusion MRI may be more sensitive to early changes.

**METHODS.:** We examined grey matter microstructure in 197 participants across the Lewy body diseases spectrum: 88 Lewy body dementia (LBD), 109 Parkinson’s normal cognition (PD-NC), and 46 controls. We compared tissue-weighted neurite density (tNDI), orientation dispersion index (tODI), and free water fraction (FWF) cross-sectionally and longitudinally. We examined relationships between microstructure, disease severity and plasma p-tau217.

**RESULTS.:** LBD participants showed widespread cortical and subcortical tODI and FWF increases compared to controls and PD-NC, despite minimal atrophy. These changes correlated with cognitive (but not motor severity) and with p-tau217. In 100 Parkinson’s patients followed-up over 3 years, cortical FWF increased in those developing cognitive decline.

**DISCUSSION.:** Microstructural measures capture early grey matter changes, offering a sensitive marker of grey matter integrity in Lewy body diseases.

## Background

Lewy body diseases, including Parkinson’s disease (PD), Parkinson’s disease dementia (PDD) and dementia with Lewy bodies (DLB) are common neurodegenerative diseases sharing overlapping clinical features and characterised by pathological accumulation of alpha-synuclein forming Lewy bodies and neurites within cortical and subcortical regions. Although there is considerable inter-individual heterogeneity in the distribution of alpha-synuclein accumulation within the brain, with different subtypes and timelines of spread proposed(1–3), current neuropathological diagnostic criteria rely on identification of Lewy pathology within the grey matter(4) and there is evidence of widespread molecular(5,6), mitochondrial(7) and functional neuronal dysfunction(8). As well as Lewy-related pathology, beta-amyloid and tau co-pathology is frequently found in LBD at post mortem, and relates to cognitive severity(9).

Conventional structural MRI measures of grey matter integrity in living patients have shown heterogeneous findings. Variable cortical atrophy patterns are seen in patients with Lewy body diseases compared to controls(10–14). Although higher rates of global cortical atrophy are associated with worse cognitive performance(12,15,16), grey matter atrophy is less reliable at earlier stages(14) and in PD patients with intact cognition(17). This suggests that measures of grey matter atrophy alone may be insensitive to early pathology, limiting their utility as biomarkers for early detection of cortical involvement.

Recent advances in diffusion-weighted imaging (DWI) have allowed examination of grey matter microstructure in-vivo. Neurite orientation dispersion and density imaging (NODDI) is a biophysical model that separates signal originating from the intra-neurite, extra-neurite and free-water compartments(18). Although mostly applied to white matter imaging, NODDI is particularly well-suited for microstructural modelling of grey matter, which mostly comprises of dendrites and crossing axons rather than long-range fibre tracts. Traditional diffusion metrics are more insensitive or ambiguous within such regions of varied orientations(18–20).

NODDI characterises three grey matter microstructural properties: 1) free-water fraction (FWF), which estimates the volume fraction of free water, contributed primarily from cerebrospinal fluid (CSF), with higher values representing increased extracellular component 2) orientation dispersion index (ODI) reflecting the spatial configuration of neurites, with higher values reflecting more complex dendritic organisation(18,20) and 3) neurite density index (NDI), reflecting the volume fraction of neurites within the brain tissue with higher values representing higher neurite density.

Such metrics of grey matter microstructure have shown increased sensitivity in capturing grey matter changes associated with healthy aging(21–23), and Alzheimer’s disease pathology(22,24–27) even in pre-symptomatic stages(21,27,28). In Lewy body diseases, few studies have comprehensively assessed grey matter microstructure across the spectrum and course of the disease. Small studies examining changes in PD patients compared to controls have consistently shown increased FWF in the basal ganglia and widespread cortical regions(17,29,30), and to lesser extent and less consistently reductions in NDI and ODI(31,32). In patients with DLB, emerging work has shown widespread changes in cortical grey matter compared to controls, with microstructural alterations correlated with beta-amyloid and tau-PET burden(33). However, PD and DLB have mostly been investigated separately with different methodologies and across different subgroups of the disease (*Table 1*).

Critically, no study has yet comprehensively characterised cortical and subcortical grey matter microstructure using NODDI across the entire continuum of Lewy body diseases or systematically examined how these metrics relate to clinical disease severity, Alzheimer’s co-pathology and longitudinal progression. Preliminary evidence suggests that progressive increases in FWF may track disease progression over time, particularly for motor symptoms(31,34) but this is less clear for cognitive progression.

In this study, we comprehensively examine microstructural properties (FWF, NDI, ODI) compared to macrostructural atrophy of cortical and subcortical grey matter across the spectrum of Lewy body diseases and examine their relationship with disease severity, Alzheimer’s co-pathology, and clinical progression. First, we examine between-group differences between patients with dementia (LBD), PD with intact cognition (PD-NC) and age-matched controls. Next, we evaluate associations with motor and cognitive disease severity and plasma p-tau217 as a marker of Alzheimer’s co-pathology. Finally, we assess whether NODDI-derived measures show longitudinal change in a subset of baseline cognitively-unimpaired PD patients followed up over 3 years.

## Methods

### Study participants

All study participants were recruited to University College London (UCL) and provided written informed consent. Ethical approval for the study was granted by the Queen Square Ethics Committee (15.LO.0476); the study was performed in accordance with the ethical standards described in the 1964 Declaration of Helsinki.

All study participants were required to be over 50 years of age and capable of informed consent. Additionally, we required for participants:

- *with Parkinson’s disease (PD):* a clinical diagnosis of PD in accordance with the Movement Disorder’s Society (MDS) clinical diagnostic criteria(35) within the past 10 years.
- *with Dementia with Lewy Bodies (DLB):* a clinical diagnosis of probable DLB as per the international consensus McKeith criteria(36).
- *with Parkinson’s Dementia (PDD): a* clinical diagnosis of Parkinson’s dementia as per MDS criteria(37), or a clinical diagnosis of PD according to MDS clinical diagnostic criteria plus both impaired function in activities of daily living due to cognitive impairment and a Montreal Cognitive Assessment (MoCA) below 26(37).
- *with Parkinson’s mild cognitive impairment (PD-MCI):* a clinical diagnosis of PD according to MDS criteria and persistent performance below 1.5 standard deviations in one test of at least two cognitive domains or in two tests in one cognitive domain(38).
- *with Mild cognitive impairment with Lewy Bodies (MCI-LB):* a diagnosis of probable MCI-LB according to consensus criteria for prodromal DLB(39).
- *Control participants:* age between 50 and 81 years and without cognitive impairment (mini-mental state examination (MMSE) greater than or equal to 25).

Participants were excluded if any confounding neurological or psychiatric disorders or contraindications to MRI were present.

Longitudinal imaging and clinical data were available in a subset of PD (n=100) and control (n=27) participants. Participants were followed-up with the same imaging and clinical protocol at 18- and 36 months(40). PD participants developing dementia or MCI at any point during follow-up were classified as having poor cognitive outcomes.

Where longitudinal imaging data were available, the latest available imaging data were used and the latest available concurrent clinical data were used to classify PD participants for cross-sectional analyses. Participants with a diagnosis of DLB, PDD, PD-MCI and MCI-LB were grouped as Lewy Body Dementia (LBD), with the remaining PD participants classified as PD-normal cognition (PD-NC).

### Clinical assessments

The detailed study protocol has been previously described in detail(40–42). In brief, global disease burden was assessed using the MDS Unified Parkinson’s Disease Rating Scale (UPRDS) total score(43), with UPDRS part III used to assess motor severity, as well as the Timed Up and Go (TUG) test(44). Cognition was assessed with MMSE(45) and MoCA(46) used as global measures of cognition and additionally with two tests per cognitive domain, including *<u>Attention</u>* (Stroop colour naming(47) and Digit Span Backwards(48)), *<u>Executive function</u>* (Category Fluency(49) and Stroop interference(47)), *<u>Language</u>* (Letter fluency(49), and Graded Naming task(50)), *<u>Memory</u>* (Word recognition(50) and Logical Memory delayed(48)) and *<u>Visuospatial function</u>* (Hooper Test of Visual Organisation (51) and Benton Judgement of Line Orientation(52)). A composite cognitive score was also calculated per participant as the averaged z-score of the MoCA plus one task per cognitive domain, as in our previous work(41). Cognitive fluctuations were assessed using the Clinician assessment of fluctuations (CAF) and One Day Fluctuations scale(53). Impairment in daily functioning was assessed using the Functional Activities Questionnaire(54), sleep using the Rapid Eye Movement Behaviour Disorder Sleep Questionnaire (RBDSQ)(55) and visual hallucinations using the University of Miami PD Hallucinations Questionnaire (UMPDHQ) (56). Levodopa equivalent daily doses (LEDD) were calculated for all PD-NC and LBD participants(57).

### Plasma collection and processing

Participants had approximately 30ml of blood collected in polypropylene EDTA tubes. Samples were centrifuged and stored at -80°C. P-tau217 concentration was measured using the AlzPath Simoa HD-X p-Tau217 Advantage-PLUS kit. The technicians performing p-tau217 measurements were blinded to clinical data. All measurements were performed in a single batch of reagents and were above the limit detection of the assay. 155 participants had a sample available for inclusion in p-tau217 analyses (66 LBD, 62 PD-NC and 27 HC).

### Image acquisition and preprocessing

All data were acquired at the same 3T Siemens Prisma scanner with the same acquisition parameters. Diffusion weighted imaging (DWI) was acquired with the following parameters: b0 (AP and PA direction), *b*=50 s/mm^2^ 17 directions, *b*=300 s/mm^2^ 8 directions, *b*=1000 s/mm^2^ 64 directions, b=2000 s/mm^2^ 64 directions, 2×2×2 mm isotropic voxels, TR=3260 ms, TE=58 ms, 72 slices, acceleration factor =2. 3D MPRAGE (magnetization prepared rapid acquisition gradient echo) was acquired with parameters: TE=3.34 ms, TR=2530 ms, TI=1100 ms, flip angle=7°, 1×1×1 mm isotropic voxels, acceleration factor=2.

Raw image data for both modalities were visually inspected for artefacts, by raters blinded to clinical data. Scans with presence of artefacts in 15 or more volumes were excluded. DWI images were pre-processed using the standard pipeline implemented in MRtrix3.0(58) including denoising(59), removal of Gibbs artefacts(60), eddy-current and motion correction(61) and bias field correction(62). Raw T1-weighted images were co-registered to the corresponding DWI image using rigid and affine registration as implemented in NiftyReg(63). A brain mask was calculated using Synthstrip(64) and transformed to DWI space. For cortical thickness calculations, the FreeSurfer v6.0 pipeline was used with default parameters.

### NODDI calculation

The NODDI model was fitted using the accelerated microstructure imaging via convex optimization (AMICO) implementation in Python(65) with a grey matter specific diffusion parameter of 1.3 μm/ms within the previously calculated brain mask, to produce voxel-wise maps for FWF, ODI and NDI. To avoid biased estimates in the presence of CSF partial volume, we calculated tissue-weighted maps (tODI and tNDI) by voxel-wise multiplying each map with the tissue fraction (66). Voxel-wise tissue-weighted tODI and tNDI maps, free-water fraction maps and cortical thickness maps were then projected to an FSLR32k surface using an algorithm weighted towards the cortical mid-thickness as previously described(20), to enable surface-based comparisons between groups.

### Statistical analysis

Demographics and clinical characteristics were compared between groups using ANOVA for normally distributed continuous variables and Kruskal Wallis for non-normally distributed continuous variables (post-hoc Tukey and Dunn respectively); the Shapiro-Wilks test and visual inspection were used to assess normality. Repeated measures ANOVA were used to assess group differences longitudinally.

Surface maps (FWF, tODI, tNDI, cortical thickness) were averaged across 200 cortical regions of the Schaeffer parcellation(67) and 32 subcortical regions of the Tian parcellation(68) and a mean value per region was calculated. For tODI and tNDI, these were weighted by the regional mean tissue fraction to minimise bias from CSF contamination or atrophy(66). Cross-sectional group comparisons (between LBD vs HC, PD-NC vs HC and LBD vs PD-NC) were performed using general linear models with age and sex as covariates and false discovery rate (FDR) correction for multiple comparisons, as implemented in BrainStat(69) for cortical regions and statsmodels for subcortical regions.

Longitudinal comparisons were performed between PD good and PD poor cognitive outcomes using general linear models with baseline age, sex and time between scans as covariates and the time * group interaction as the variable of interest, FDR correcting for multiple comparisons.

Correlations with metrics of disease severity were also performed using the same general linear models and covariates across all disease participants (excluding controls), correcting for age and sex and FDR-correcting for multiple comparisons.

### Availability of data and materials

All processing and analysis code is made available through: https://github.com/AngelikaZa/GM_Microstructure

Anonymised group level data are also available through the same repository. Individual level data and raw imaging data may be shared upon reasonable request to the corresponding author.

## Results

We included 243 participants: 88 LBD (34 DLB, 34 PDD, 17 PD-MCI, 3 MCI-LB), 109 PD-NC and 46 controls. LBD participants were older, had a higher proportion of males and had worse cognition (lower composite cognitive scores, MoCA and MMSE), higher overall disease burden and higher rates of hallucinations, cognitive fluctuations and REM sleep behaviour disorder. Demographic and clinical information for our cohort is seen in *Table 2*.

LBD subgroups did not significantly differ in any demographic or clinical characteristics, other than lower composite cognitive scores in DLB participants and higher overall disease duration in PD-MCI and PDD participants, as expected (see *Table 3* for details).

A subset of 100 PD participants and 27 controls underwent longitudinal clinical and imaging assessments at 18- (Session 1) and 36-months (Session 2). All PD participants were cognitively intact at their baseline visit but 31 developed MCI or dementia during follow-up (PD poor outcomes). Demographics and clinical characteristics of our longitudinal cohort have been previously described in detail(40); baseline between group differences are also presented in *Table 4*.

### Widespread cortical and subcortical grey matter microstructural alterations are seen in the absence of atrophy, in LBD participants compared to controls and PD-NC

LBD participants showed widespread changes in grey matter microstructure compared to controls and to a lesser extent compared to PD-NC (general linear models across 200 cortical and 32 subcortical regions, with age and sex as nuisance covariates, FDR-corrected for multiple comparisons). LBD participants showed widespread increases in FWF in both cortical (*Figure 1A*) and subcortical regions (*Figure 1B*) compared to controls. They also showed increases in FWF compared to PD-NC primarily involving pre-frontal, occipital and medial temporal regions as well as hippocampi bilaterally.

**Figure 1.**
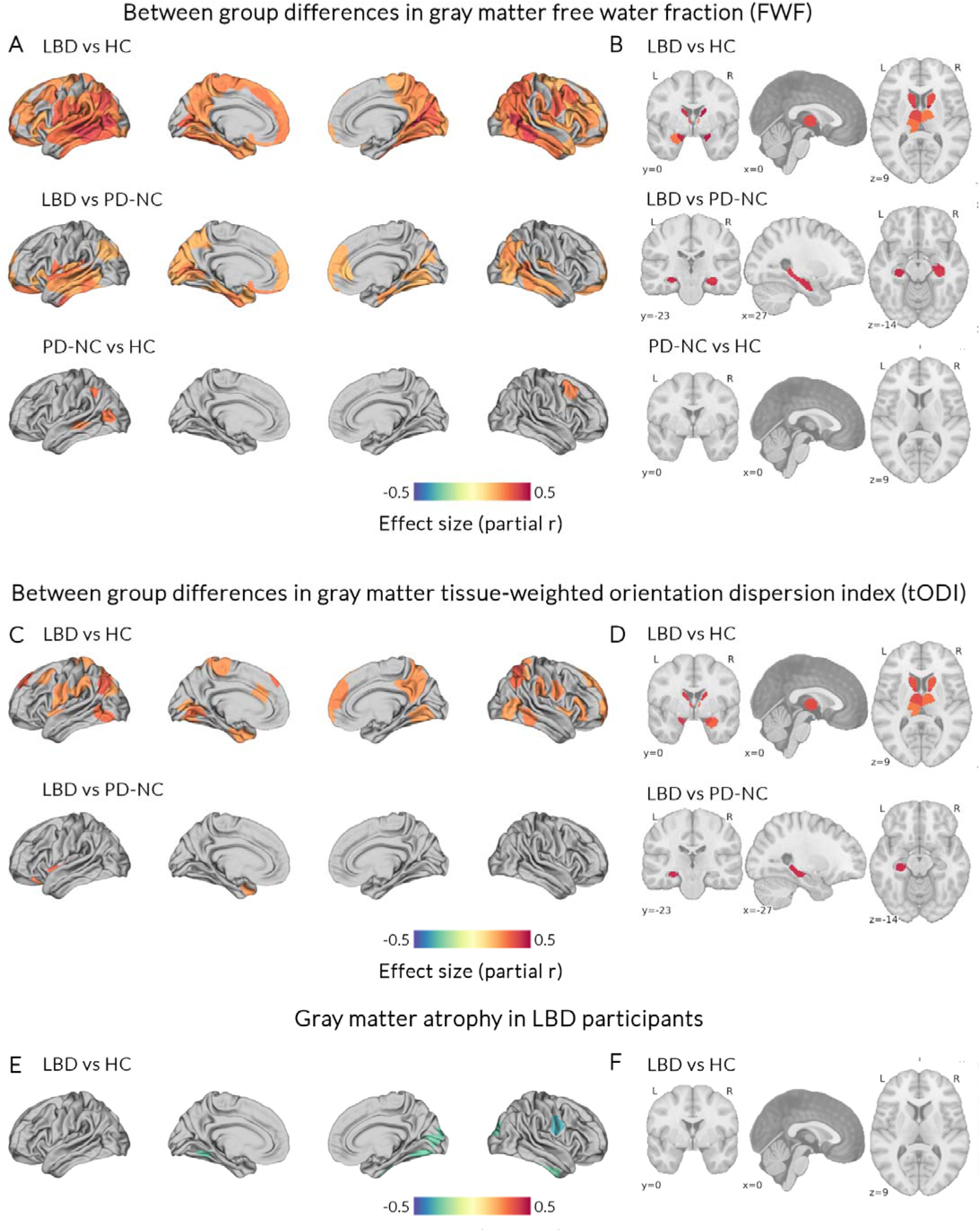
Microstructural grey matter changes across the spectrum of Lewy body diseases. Between group differences were examined using general linear models with age and sex as nuisance covariates and FDR-correction for multiple comparisons, in 197 participants across the spectrum of Lewy body diseases and 46 controls. We compared: LBD (n=88) vs HC (n=46), LBD vs PD-NC (n=109) and PD-NC vs HC across 200 cortical and 32 subcortical regions . **Free-water fraction (FWF)** compared between groups in A) cortical and B) subcortical regions. **Tissue-weighted orientation dispersion index** (tODI) compared between groups in cortical (**C**) and subcortical regions (**D**). **Cortical thickness** (**E**) and **subcortical grey matter volume** (**F**) compared between groups. Colour scale depicts effect size (blue colours decreases in the examined group; red colours increases). Only statistically significant regions after multiple comparisons correction (FDR-p <0.05) are shown. There were no statistically significant group differences in tissue-weighted neurite density index. LBD: Lewy body dementia patients; PD-NC: Parkinson’s disease patients with normal cognition; HC: Age-matched controls; FWF free water fraction; tODI: tissue weighted-orientation dispersion index; FDR false discovery rate.

LBD participants also showed increases in tODI reflecting increased complexity in dendritic orientation, compared to controls; this had similar regional distribution as the increases seen in FWF but involved less widely distributed regions, with tODI changes primarily seen within the parietal, temporal and frontal lobes (*Figure 1C*) and basal ganglia (*Figure 1D*). Increases in tODI were also seen in LBD compared to PD-NC within right frontal and temporal regions and the right hippocampus.

There were no changes in tNDI for any of the group comparisons in any cortical or subcortical regions. Additionally, despite widespread changes in FWF and tODI, there was limited cortical atrophy (Figure 1E) and no statistically significant subcortical atrophy in LBD compared to controls (Figure 1F); there were no changes between LBD and PD-NC in either cortical or subcortical regions.

In contrast to the changes seen in LBD participants, there were very limited changes in PD-NC participants compared to controls, with increases in FWF alone in left temporo-parietal and right frontal regions only and no changes in PD-NC compared to controls in tODI (*Figure 1A*).

There were no statistically significant differences in any metrics between any of the LBD subgroups (DLB vs PDD). Full subcortical region of interest results are presented in *Supplementary Tables 1, 2 and 3*.

### Microstructural alterations are associated with cognitive but not motor disease severity and with plasma p-tau217 levels

To examine whether these changes were relevant to disease severity across the spectrum of Lewy body disease, we examined the association between mean regional FWF and tODI and different clinical severity metrics of cognition (composite cognitive score, MOCA and MMSE), overall disease burden (total UPDRS score) and motor severity (UPDRS part 3, TUG score) across all disease participants, including both LBD and PD-NC. Increased cortical FWF and to a lesser extent tODI signal was significantly correlated with worse cognitive performance (composite cognitive score presented in *Figure 2A*, with similar changes seen in relation to MOCA and MMSE, seen in *Supplementary Figure 1*). Increased cortical FWF was also correlated with higher overall disease burden (*Figure 2B*). In contrast, cortical thickness was not significantly correlated with cognitive or overall disease severity. There were no significant correlations with cortical FWF, tODI signal or thickness with any motor severity metric.

**Figure 2.**
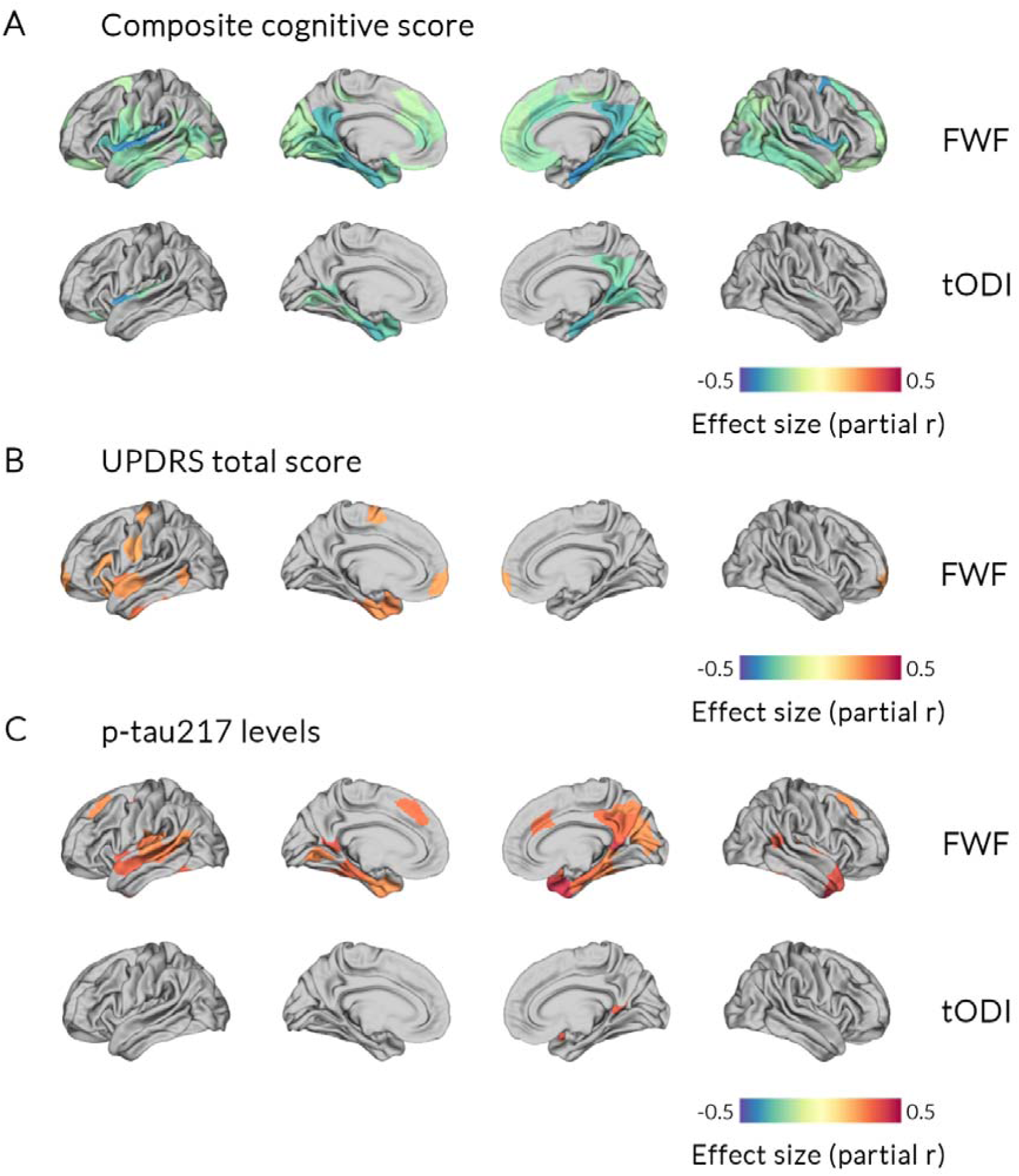
Microstructural grey matter alterations in Lewy body disease are correlated with cognitive but not motor symptoms and with amyloid co-pathology. **A.** Increased cortical free-water fraction (FWF) and tissue-weighted orientation dispersion index (tODI) were correlated with lower composite cognitive scores in patients with Lewy body diseases. **B.** Increased FWF alone within prefrontal, temporal and parietal regions was associated with global disease severity in patients with Lewy body disease, measured using the Movement Disorders Society Unified Parkinson’s disease rating scale (UPDRS total score). **C**: Increased FWF within temporal and parietal regions and increased tODI within the left temporal lobe were associated with increased plasma p-tau217 levels, a marker of overall amyloid and tau co-pathology burden. General linear models were used with age and sex as nuisance covariates and false discovery rate (FDR) correction for multiple comparisons across all participants with Lewy body diseases (n=155). Colour scale depicts effect size (blue colours decreases in the examined group; red colours increases). Only statistically significant regions after multiple comparisons correction (FDR-p <0.05) are shown.

For subcortical regions, only mean right hippocampus tODI signal was correlated with overall disease severity (total UPDRS score) with no other significant correlations for subcortical FWF and tODI signal and any disease severity metrics.

Regional FWF and tODI cortical, but not subcortical signal was also correlated with plasma p-tau217 levels, particularly within temporo-parietal regions (*Figure 2C*). There were no correlations between tNDI and clinical or plasma measures.

### In Parkinson’s patients, longitudinal microstructural grey matter changes track poor cognitive outcomes

Finally, we assessed whether measures of grey matter microstructural integrity track disease progression longitudinally. To do this, we examined longitudinal group*time interaction in a subset of 100 PD participants followed up over 3 years. We compared how slopes of change differed between those participants who progressed to PD-MCI or dementia (PD poor outcomes (n=31)) to those whose cognition remained stable (PD good outcomes (n=69)).

There were higher longitudinal increases in FWF in PD poor outcomes compared to PD good outcomes within temporal regions (*Figure 3A*), right thalamus and right caudate (*Figure 3B*). There were no significant differences in rate of change of tODI or tNDI between PD poor and PD good outcomes. FWF rate of change within the right limbic temporal pole was significantly correlated with longitudinal rate of change in composite cognitive scores (*Figure 3C*) and MOCA (*Supplementary Figure 2*) but not with motor or overall disease severity.

**Figure 3.**
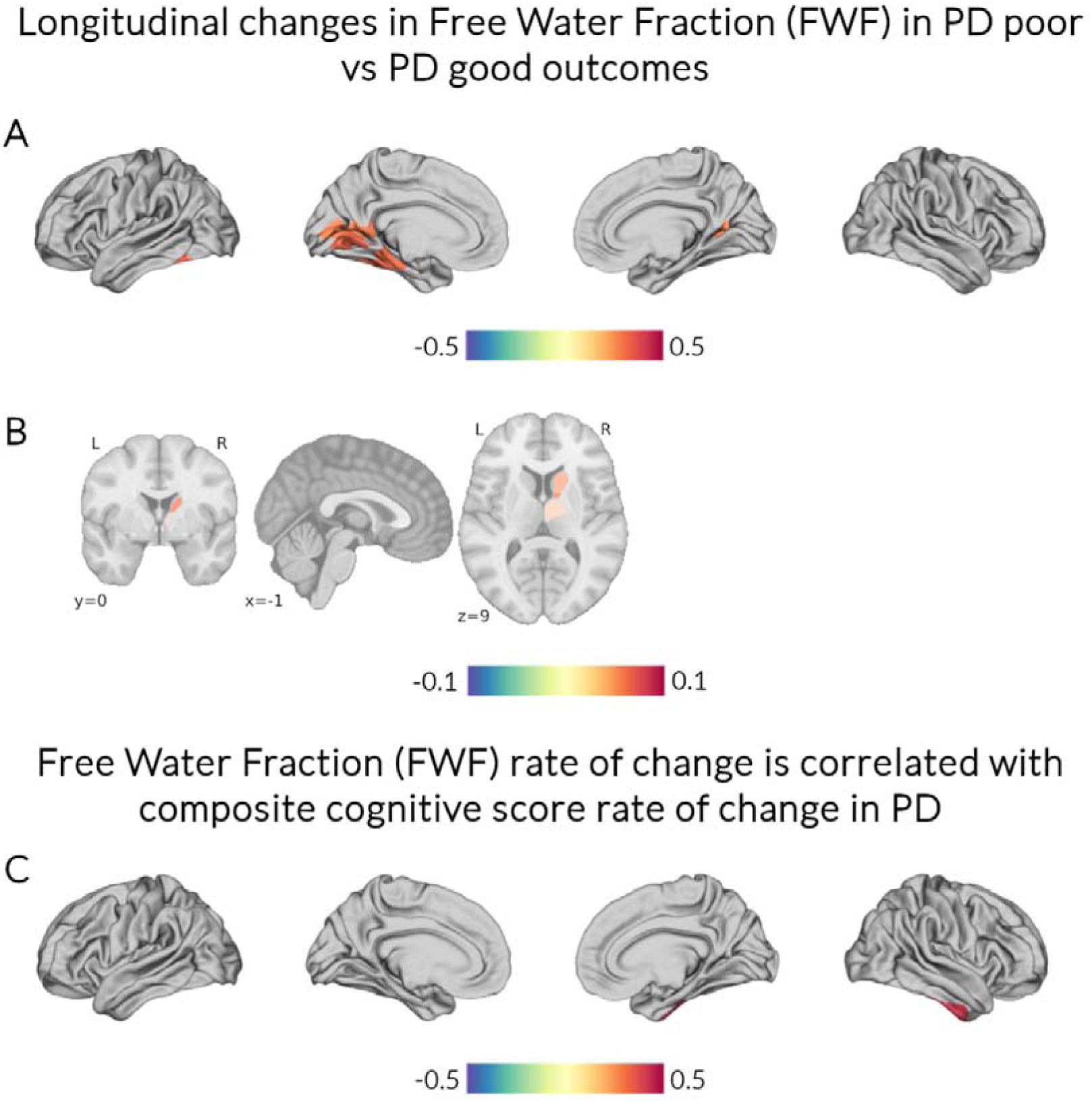
Free water fraction tracks with longitudinal cognitive decline in Parkinson’s disease. General linear models were used to assess longitudinal differences in group * time interactions between patients with Parkinson’s disease that developed mild cognitive impairment or dementia (PD poor outcomes, n=31) compared to those with stable cognition (PD good outcomes, n=69); age and sex as nuisance covariates, false discovery rate (FDR) correcting for multiple comparisons. **Larger increases in free water fraction (FWF) were seen longitudinally in PD poor compared to PD good outcomes** in **A)** cortical regions (temporal and occipital regions) and **B)** subcortical regions (right basal ganglia and thalamus). **C) Rate of change in FWF was correlated with rate of change in composite cognitive scores** within the right limbic temporal pole. Colour scale depicts effect size (blue colours decreases in the examined group; red colours increases). Only statistically significant regions after multiple comparisons correction (FDR-p <0.05) are shown.

There were no significant differences in longitudinal cortical thickness change between PD poor and PD good outcomes.

## Discussion

Here we comprehensively examine grey matter microstructural integrity in 243 participants spanning the full Lewy body disease spectrum. We show widespread grey matter cortical and subcortical microstructural alterations in the absence of significant cortical atrophy. LBD participants showed increased FWF and tODI compared to both controls and PD with intact cognition, with changes correlating with cognitive but not motor severity. FWF and tODI also correlated with Alzheimer’s co-pathology burden measured using plasma p-tau217.

Longitudinally, PD participants who later developed cognitive impairment showed faster increases in FWF within temporal and subcortical regions, correlating with cognitive decline. Together, these findings indicate that microstructural grey-matter changes, particularly elevated FWF, reflect early, progressive grey matter pathology related to cognitive deterioration and tau co-pathology in Lewy body disease.

These measures reflect different microstructural properties. FWF refers to the fraction of the diffusion signal attributable to extracellular water, which is not restricted by cell membranes or neurite structure (for example CSF or enlarged interstitial space)(18). Therefore, an increase in FWF may reflect extracellular fluid accumulation (e.g., vasogenic oedema), blood-brain barrier disruption, tissue atrophy below the threshold observable as gross structural changes, or neuroinflammation. tODI refers to the degree of angular variation of neurites within a voxel, with increases suggesting more complex orientations, for example increased dendrite branching(18,70). Finally, tNDI estimates the volume fraction of intra-neurite water and is therefore a proxy for neurite density, with lower tNDI reflecting neurite loss(18,24).

Relatively few studies have assessed grey matter microstructure, particularly within the cortex, in Lewy body diseases (for summary, see *Table 1*). Increased free-water fraction has been shown separately in patients with Parkinson’s disease(17) and DLB(33,34) compared to controls. In these studies, free-water fraction was mainly reported in relation to measures of motor severity(17,31,33,34). In keeping with these previous studies, we found widespread increases in FWF in patients with Lew body dementia compared to controls and to those with Parkinson’s with intact cognition. Our study extends previous work by including all clinical stages of Lewy body diseases in one clinical cohort. We show that frontal, temporal and occipital changes in FWF correlate specifically with cognitive but not motor severity measures. Additionally, we show that FWF within temporal regions can track longitudinal cognitive impairment in Parkinson’s disease, providing further evidence for microstructural grey matter alterations specifically accompanying cognitive decline.

In addition, we found increased tODI in LBD compared to controls and to a lesser extent to PD-NC, which again was correlated with cognitive severity but was not associated with longitudinal progression. Mak et al. recently described modest reductions in ODI compared to controls(33), with reductions in the right frontal lobe and caudate also described in a small study of PD-MCI compared to PD-NC(29). However, both these studies had smaller sample sizes, did not employ tissue-correction for ODI and, crucially, had a more severely affected cohort from a cognitive perspective with a mean 3 points additional MMSE reduction described in both cohorts compared to ours(29,33). The differences in our findings could reflect differences in disease stage. NODDI-derived tODI strongly correlates with dispersion in histological validation studies(71,72) and within the grey matter dispersion mainly reflects sprawling dendritic processes and to a lesser extent axonal dispersion. Axonal and dendritic structural dysfunction has been implicated as an early process in alpha-synucleinopathies, occurring before overt axonal or neuronal loss: exogenous alpha-synuclein in human and animal cell cultures induces dendritic spine impairment leading to dysmorphic dendrites and axons(73–75), overexpression leads to axonal swelling(76), and increases in spine density are seen early within the prefrontal cortex of transgenic mice(77). This increased dendritic beading, axonal swelling, and dendritic cytoskeletal disorganisation early in the course of the disease would lead increase tODI values at early stages; our study provides additional evidence that this may be captured in-vivo.

In contrast to the widespread changes in FWF and tODI in LBD participants, we found no significant differences in tNDI across any group, consistent with the minimal atrophy also observed in our cohort. Whilst inevitably neuronal loss occurs in Lewy body diseases, preserved neuronal density early in the course of the disease is consistently seen following exogenous alpha-synuclein seeding in mice despite evident neuronal functional alterations(74,78,79). Our findings, together with previous neuroimaging studies showing high heterogeneity in measures of grey matter atrophy (10–14,17,80) further support that metrics reflecting overt neuronal loss are less sensitive as a biomarker in Lewy body diseases, particularly in earlier stages.

This is in contrast to changes seen in Alzheimer’s disease, where reductions in cortical neurite density and reduced orientation dispersion (reduced NDI and ODI) are consistently described, especially in temporo-parietal and medial temporal cortices(22,25,26,81).

Decreases in temporal NDI and ODI are seen along the MCI to Alzheimer’s disease continuum(21,22,24,28), and are linked to both memory decline and tau and amyloid PET burden(22,25–28). This suggests that progressive loss of neurite density and complexity is a core cortical signature of Alzheimer’s pathology. In contrast, we have shown a divergent NODDI profile in Lewy body diseases, with prominent FWF increases in cortical and subcortical grey matter, preserved NDI and locally increased tODI early in the disease course. Importantly, however, tODI may be preserved or reduced in later stages(17,33). This divergent suggests that NODDI-derived measures may be useful as potential diagnostic biomarkers as well differentiating DLB from Alzheimer’s disease; this could be particularly relevant given the high prevalence of amyloid co-pathology in DLB(9) which could lead to an Alzheimer’s misdiagnosis if only CSF/plasma amyloid biomarkers are considered.

We did not find widespread group differences for cortical atrophy or associations between cognitive severity and cortical atrophy using conventional atrophy measures. This is consistent with previous work showing variability in cortical atrophy even between LBD and PD-NC(15,80); and for associations with cognition(10,12–15). For PD, cortical atrophy is even more inconsistent, with many studies showing little or no atrophy compared with controls, and with cognitive severity(15,17,30,82). Our finding of consistent group differences and associations between FWF and cognitive severity cross-sectionally and over time, suggests that this measure is more sensitive to relevant tissue changes. Interestingly, FWF seems to also be sensitive to different tissue changes than quantitative susceptibility mapping (QSM), which is most sensitive to iron accumulation. We have previously shown differences in QSM between PDD and DLB and QSM sensitivity to motor severity(83). In contrast, FWF appears more sensitive to cognitive rather than motor severity and more sensitive than QSM to longitudinal changes in PD(84). As FWF reflects higher levels of free water this may reflect changes such as extracellular fluid expansion, oedema, disruption to tissue architecture or neuroinflammation(85,86).

In our cohort, we further observed that higher FWF and tODI within temporal and parietal regions correlated with plasma p-tau217 levels, but this did not account for the full microstructural correlations seen in LBD patients compared to cognitively intact patients and controls. Whilst amyloid co-pathology in Lewy body diseases is widespread, tau co-pathology is generally localised within the temporal neocortex(9,87). This corresponds with the regions showing correlations with p-tau217 levels in our cohort. Increased tau burden in patients with DLB has been linked with worse cognitive severity in-vivo(42,88) and with greater alpha-synuclein cortical load at post-mortem(87). Our findings therefore suggest that microstructural alterations observed in alpha-synucleinopathies could reflect combined effects from both alpha-synuclein and amyloid and tau co-pathology.

Our study has certain limitations. Ours was a single centre study and longitudinal data were available only on a subset of Parkinson’s patients. Although our cohort was enriched for cognitive impairment with 31 participants developing dementia or MCI, larger multi-centre longitudinal studies could validate the utility of FWF as a biomarker that can track cognitive disease progression in Lewy body diseases. Additionally, although our observed alterations in NODDI-derived metrics in Lewy body dementia patients show distinct patterns from those described in Alzheimer’s disease, did not directly compare these groups. Future work could explore the role of NODDI-derived metrics as a biomarker supporting in-vivo differential diagnosis. Finally, although we examine the effect of global amyloid and tau co-pathology in microstructural grey matter alterations through plasma p-tau217 levels, we did not have measures of abnormal alpha-synuclein burden in our cohort. Future research incorporating more specific measures of pathological protein accumulation such as amyloid PET, tau PET and alpha-synuclein seed aggregation assays could disentangle the causes of grey matter microstructural change in Lewy body diseases.

In summary, we comprehensively assess for the first time grey matter microstructure and macrostructure across the whole spectrum of Lewy body disease cross-sectionally and longitudinally. We show widespread increases in cortical and subcortical free-water fraction and elevated tissue-weighted orientation dispersion in the absence of significant atrophy.

This indicates extracellular expansion and dendritic disorganisation rather than overt neuronal loss. These changes were specific to cognitive and not motor disease severity, and related to plasma p-tau217. We also showed that free-water fraction increased over time in Parkinson’s patients who developed cognitive decline during follow-up. Together, our results suggest that NODDI-derived measures, particularly free-water fraction, capture early and evolving grey matter changes in Lewy body diseases and could be a sensitive marker for disease staging and longitudinal tracking of cortical integrity.

